# Engineering 3D Skeletal Muscle Tissue with Complex Multipennate Myofiber Architectures

**DOI:** 10.1101/2025.01.06.631119

**Authors:** Maria A. Stang, Andrew Lee, Jacqueline M. Bliley, Brian D. Coffin, Saigopalakrishna S. Yerneni, Phil G. Campbell, Adam W. Feinberg

**Affiliations:** Department of Materials Science & Engineering, Carnegie Mellon University, Pittsburgh, PA 15213, USA; Department of Biomedical Engineering, Carnegie Mellon University, Pittsburgh, PA 15213, USA

**Keywords:** 3D bioprinting, skeletal muscle, tissue engineering, FRESH, collagen

## Abstract

The hierarchical architecture of skeletal muscle spans from microscale sarcomeres to macroscale myofibers and is integral to its contractile functionality. Pathologies such as volumetric muscle loss (VML) compromise this structure and destroy the native extracellular matrix (ECM), exceeding the regenerative capacity of endogenous repair mechanisms. Here, we present a novel method for tissue engineering biomimetic three-dimensional (3D) skeletal muscle with complex architectures by leveraging freeform reversible embedding of suspended hydrogels (FRESH) 3D bioprinting of the ECM. Collagen type I scaffolds mimicking diverse muscle architectures— including parallel, unipennate, bipennate, multipennate, and convergent—were designed, FRESH printed, and seeded with C2C12 myoblasts to guide myogenesis. Engineered muscle tissues demonstrated scaffold-mediated alignment and fusion into functional myotubes, exhibiting contractile responses to electrical stimulation with architecture-dependent specific force of ∼1 kN/m^2^ and a positive force-frequency relationship. *In vivo* implantation further revealed scaffold-directed cellular and vascular organization, underscoring the translational potential of this approach. In summary, this study demonstrates the capability to use FRESH 3D bioprinting to engineer physiologically relevant muscle architectures, significantly advancing the design of functional muscle tissues for regenerative medicine and in vitro modeling applications.

## 1. Introduction

Structure-function relationships exist across every length scale in living systems, from the molecular and cellular level to the tissue and organ level and finally to the whole organism. This is evident in the hierarchical organization of skeletal muscle, the primary actuation system in the human body. The sarcomere, which is the fundamental repeat unit of the contractile machinery, is made up of overlapping arrays of actin and myosin filaments. At the same time, muscle fibers and fascicles are oriented in a range of architectures that impact 3D muscle deformation during contraction.^1,2^ For instance, parallel muscles are made up of long fibers with many sarcomeres in series and can thus achieve greater contractile velocities; conversely, pennate muscles are made up of shorter fibers and more sarcomeres in parallel, enabling greater force generation.^3^ Upon muscle injury or disease, the underlying structure can be damaged or lost, resulting in reduced or impaired function. Although skeletal muscle can regenerate minor injuries, loss of muscle function, known as volumetric muscle loss (VML), can result from disease (e.g., myopathies or neuromuscular disorders), traumatic injuries, or surgical resections.^4,5^ This irreversible damage can be attributed to the destruction of the basal lamina, which houses muscle satellite (progenitor) cells that are required for muscle regeneration.^6^ The basal lamina is also a key extracellular matrix (ECM) structure needed for myogenesis as it provides instructive cues to satellite cells and their differentiated myoblast progeny to fuse and align into organized and functional myofibers.^7^ Although there is no cure for VML, a primary therapeutic option is autologous (a.k.a., functional free) muscle transfer, in which healthy tissue is grafted from an unaffected donor site.^8^ However, full muscle function is not recovered due in large part to fibrosis, and patients face complications such as donor site morbidity, infection, and/or necrosis that can all lead to graft failure.^9^

Engineered muscle tissue is a potential solution to restore muscle function, especially in injuries where the underlying basal lamina is lost and regeneration is limited.^10,11^ The ECM is critical for muscle regeneration, and a number of tissue engineering strategies have used ECM proteins to guide myoblast fusion and myotube alignment to rebuild muscle architecture in vitro. For example, patterning of ECM proteins like laminin into microscale lines via microcontact printing has been shown to align myotubes on 2-dimensional (2D) surfaces.^12^ Similarly, muscle cells have been aligned using engineered substrate micro-topographies in the form of aligned fibers, grooves, posts or holes.^9,13,14^ This concept has also been extended towards 3-dimensional (3D) muscle tissue using an array of posts in a predetermined orientation to create thin and porous skeletal muscle tissue constructs.^15^ Other 3D muscle tissue constructs have been engineered using mechanical loading and/or electrical stimulation to achieve greater maturation and improved contractile function.^16,17^ This is often accomplished by casting and gelling a cell and ECM hydrogel mixture around two or more anchor points followed by cell-mediated compaction. The isometric forces generated drives cellular alignment along the axis of tension.^15,18,19^ While these 3D approaches have demonstrated significant progress, it has remained challenging to engineer muscle tissues with greater geometric complexity beyond muscle tissue constructs with parallel myofibers that align to the direction of tension. Current strategies have been unable to recapitulate a broad range of skeletal muscle types in the body, such as unipennate, bipennate and multipennate myofiber architectures. Muscles that have pennate architectures are comprised of myofibers that are oriented at an angle and are thus typically shorter than those found in parallel muscles. Given a constant muscle volume, pennate muscles possess a larger quantity of myofibers and therefore a greater number of myotubes in parallel, yielding a larger physiological cross-sectional area and force generation.^1–3^ There is thus a need for new tissue engineering approaches that can guide myogenesis in 3D to direct microscale myotube alignment while simultaneously recreating the macroscale native skeletal muscle architectures that direct function.

Here, we have developed a new approach to engineer 3D skeletal muscle tissues with complex myofiber architectures by controlling the spatial patterning of ECM proteins to mimic native muscle organization. To do this we used collagen type I, the primary ECM component of skeletal muscle, and built acellular scaffolds that would serve as physical guidance cues for muscle formation. This required a new biofabrication approach, for which we leveraged the advanced freeform reversible embedding of suspended hydrogels (FRESH) 3D bioprinting process that has specifically demonstrated the engineering of collagen constructs at extremely high resolution.^20,21^ Using established casting methods, we show that seeding C2C12 myoblasts on these collagen scaffolds leads to the formation of biomimetic muscle tissue constructs with structural complexity that exceeds current state-of-the-art. Specifically, FRESH was used to 3D bioprint acellular collagen type I scaffolds mimicking five skeletal muscle architectures (parallel, unipennate, bipennate, multipennate, and convergent). Engineered muscle tissues were created by casting C2C12 myoblasts in a collagen gel directly onto the scaffolds, where the FRESH printed collagen filament guided myoblast fusion and myotube alignment. C2C12 myoblasts infiltrated into the scaffold, conformally compacted around the collagen filaments, and then due to spatial confinement fused into myotubes in the defined 3D spatial pattern. Furthermore, these tissues were anchored at either end throughout the cell culture period, further enhancing cellular alignment through isometric tension. Contractile function was assessed, showing that the engineered muscle tissues were responsive to electrical stimulation, generated force, and displayed expected calcium handling. Further, subcutaneous implantation *in vivo* showed that scaffold guidance was maintained, biasing the organization of infiltrating host cells into the engineered muscle tissue. Overall, this approach shows the ability to engineer contractile muscle tissue that addresses a much broader range of anatomical architectures than previous approaches.

## 2. Results & Discussion

### 2.1. Skeletal Muscle Tissue Engineering Workflow Using FRESH 3D Bioprinting

To engineer 3D skeletal muscle tissue, we developed a multi-step approach that consists of: (i) collagen scaffold fabrication via FRESH 3D bioprinting, followed by (ii) cell casting in a collagen hydrogel solution **(Fig. 1A)**. The design of each scaffold was inspired by native muscle architectures and created through a combination of computer-aided design (CAD) and slicing software, where print parameters like infill pattern (spatial organization of filaments), density (degree of filament packing), and angle (filament orientation) were selected to mimic muscle fiber alignment, width, and pennation angle, respectively. These settings were used to slice the 3D CAD model into layers and generate the printing instructions, or G-code, for the 3D printer. Using our custom-designed bioprinters,^22,23^ acidified type I collagen was FRESH printed^20^ in the desired architecture, and each scaffold was subsequently released and transferred to a sterile PDMS chamber. These chambers contained two or more PDMS posts to act as mechanical anchor points during the cell casting and culture periods. To create tissues, C2C12 myoblasts were cast in a hydrogel solution into the chamber around the FRESH printed collagen scaffolds (**Fig. 1B**). In the absence of a scaffold to direct alignment, these casting approaches result in bundles with myotube alignment confined to one direction, parallel to the axis of tension, similar to the way many 3D linear muscle bundles are engineered.^24–27^ By comparison, our scaffold-based patterning approach directs alignment in multiple, predetermined orientations to mimic a variety of skeletal muscle architectures.

**Fig. 1.**
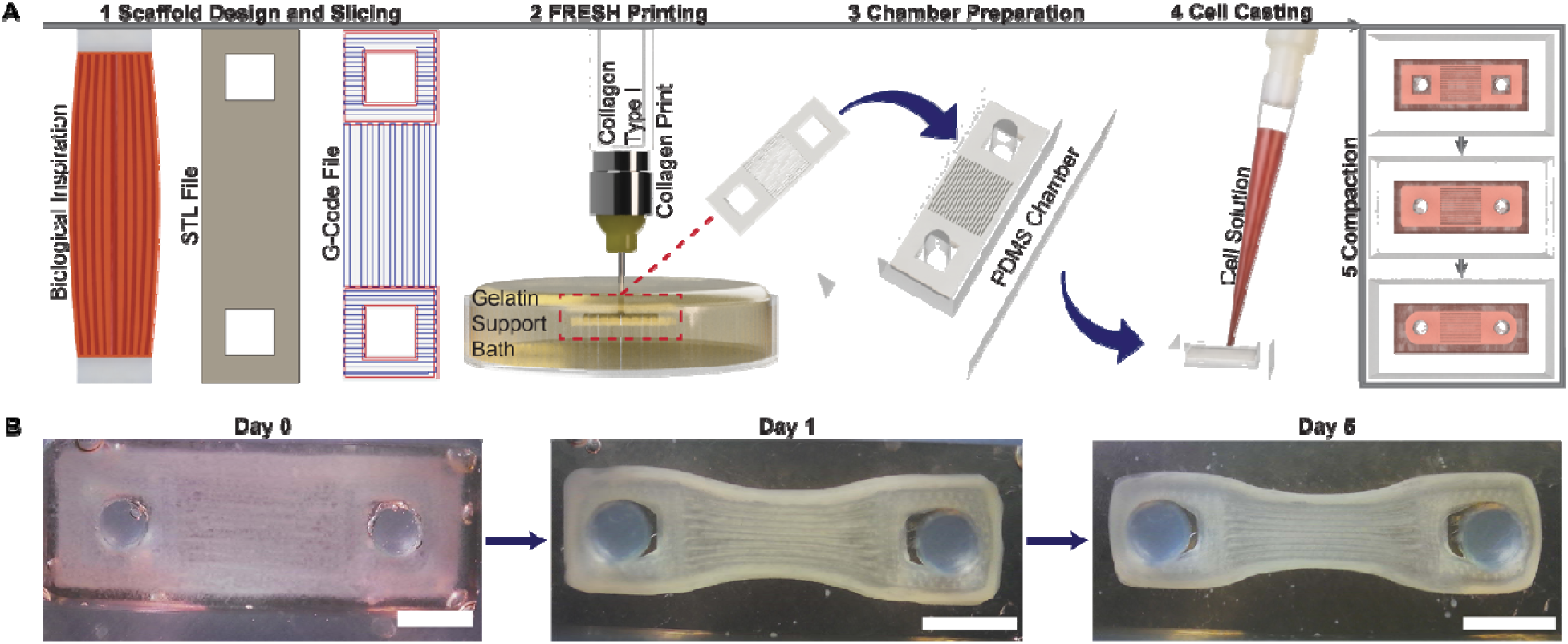
Workflow for engineering skeletal muscle tissue with biomimetic architectures. (**A**) Collagen scaffolds are designed using CAD software and converted from an STL file into G-Code using slicer software (Step 1). The collagen scaffold is FRESH 3D bioprinted, released, and washed (Step 2), and then carefully transferred to a sterile PDMS chamber (Step 3). A C2C12 cell laden collagen solution is pipetted directly onto the scaffold (Step 4), resulting in conformal compaction around it (Step 5). (**B**) Time-lapse microscopy showing that over multiple days, cell-mediated compaction around the scaffold occurs. Scale bars: 2 mm.

### 2.2. Design and FRESH 3D Bioprinting of Collagen Scaffolds with Biomimetic Muscle Architectures

To engineer biomimetic skeletal muscle tissue, three native skeletal muscle architectures – parallel, unipennate, and bipennate – were used as templates to guide the design of collagen scaffolds **(Fig. 2A)**. In muscles possessing a parallel architecture, muscle fibers and fascicles (bundles of myofibers) are arranged parallel to the longitudinal axis of the tissue, while the fascicles in pennate muscles insert obliquely to a central tendon, typically at angles ranging from 0 to 30 degrees in mammalian muscles.^28,29^ Additionally, muscle fibers can be up to 100 μm in diameter. Print parameters (e.g., infill pattern, angle, density) were selected to mimic these native muscle features to create scaffolds whose 3D filament arrangement would guide myoblast fusion and alignment in the desired configuration. To mimic the muscle fiber orientation, aligned rectilinear infill was chosen for each model, consisting of filaments parallel to one another in the designated direction. For the parallel architecture, an infill angle of 0° produced filaments oriented along the longitudinal axis of the scaffold. Similarly, unipennate and bipennate models were designed with infill angles of +30° and +30°/-30°, respectively. Finally, all scaffolds were designed with an infill density of 40%, corresponding to an extruded collagen filament diameter of ∼100 µm and spacing between filaments of 80-90 μm. Mimicking native muscle fiber widths, these print parameters were carefully chosen such that the printed collagen filaments could provide contact guidance, comparable to surface patterning and microtopography of ∼100 μm shown to be effective in aligning cells.^30^ This confines fusion and alignment to the void spaces between filaments, resulting in macroscale arrangement of highly aligned muscle fibers in the engineered geometry.

**Fig. 2.**
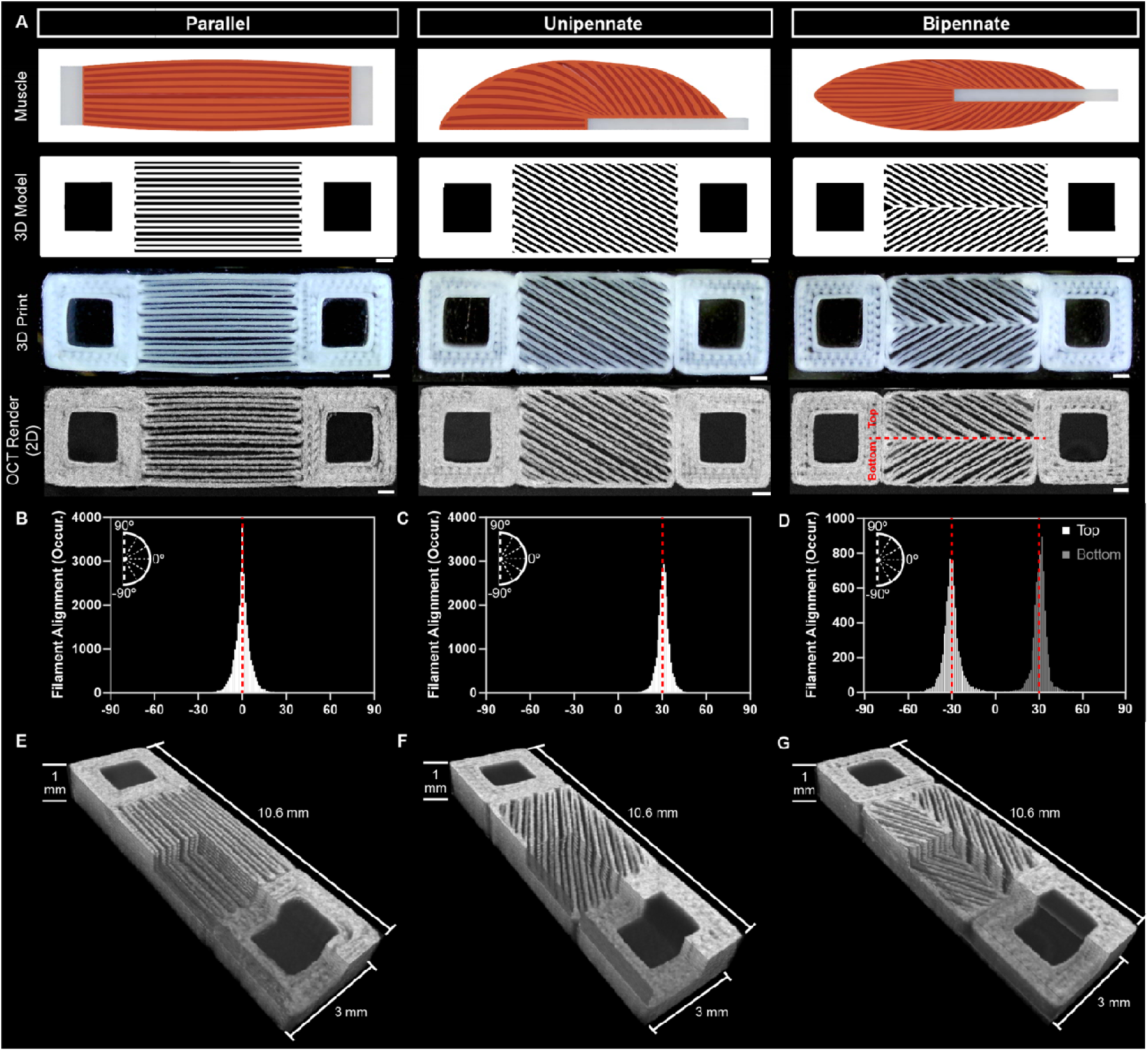
Design and filament alignment analysis of FRESH 3D bioprinted collagen scaffolds. (**A**) Collagen scaffolds recapitulating parallel, unipennate, and bipennate muscle architectures were designed in slicing software and FRESH 3D bioprinted. Printed scaffolds were imaged using OCT, and a maximum intensity Z-projection of each stack was created. Scale bars: 500 μm. (**B-D**) Histograms of collagen filament alignment for (**B**) parallel, (**C**) unipennate, and (**D**) bipennate muscle architectures. Filament alignment was quantified, where an orientation angle of 0° is designated as the longitudinal axis of the scaffold. Red dashed lines indicate the intended orientation angle from the G-Code. (**E-G**) 3D rendering of the OCT images for (**E**) parallel, (**F**) unipennate, and (**G**) bipennate muscle architectures. 3D cutouts from each scaffold demonstrate consistent filament alignment throughout the length, width, and depth.

To quantitatively assess print fidelity, optical coherence tomography (OCT) was used to image the scaffolds in 3D to determine filament orientation and pennation angle in the engineered tissue scaffolds (Fig. 2A). Collagen filaments were well aligned across all three architectures, with narrow distributions centered around the intended orientation angle (denoted by the red dashed lines, **Fig. 2B-D)**, The parallel and unipennate models were aligned at angles of 0° and 31°, respectively. The bipennate model, whose design consists of opposing pennation angles of -30°/30°, displayed two sharp peaks centered around -31°/32°. The printed constructs were thus in good agreement with the models, which was demonstrated further with 3D OCT renders **(Fig. 2E-G)**. 3D cutouts from each scaffold revealed consistent filament alignment and spacing through each construct dimension. This analysis demonstrates that the FRESH 3D bioprinting platform can be utilized to create reproducible 3D ECM scaffolds that may be suitable for the engineering of skeletal muscle tissues with a variety of pennation angles.

### 2.3. FRESH Printed Muscle Tissues Mimic Native Muscle Architectures

After validating scaffold architecture, we next investigated the ability to drive the formation of highly aligned myotubes in the engineered scaffold geometries. To assess this, C2C12 myoblasts in a collagen gel were cast around the FRESH printed scaffolds, allowed to infiltrate and compact, and then differentiated into myotubes (Fig. 1B). Immunofluorescent imaging of parallel, unipennate, and bipennate architectures revealed that the C2C12 myoblasts fused into multinucleated myotubes predominantly organized and aligned in the direction of the collagen filaments, regardless of scaffold architecture **(Fig. 3A)**. The FRESH 3D bioprinted scaffolds provided directional cues at the cellular scale, where the collagen filaments guided myogenesis through contact guidance, and organized myofibers at the macroscale to create biomimetic 3D muscle architectures. It should be noted that a small fraction of myotubes located at the surface of the tissues were not aligned to the collagen filaments, likely because they were separated from the topographical cues of the collagen scaffold by one or more layers of myotubes. This demonstrates the need for direct contact guidance of the collagen filaments in organizing myotube orientation.

**Fig. 3.**
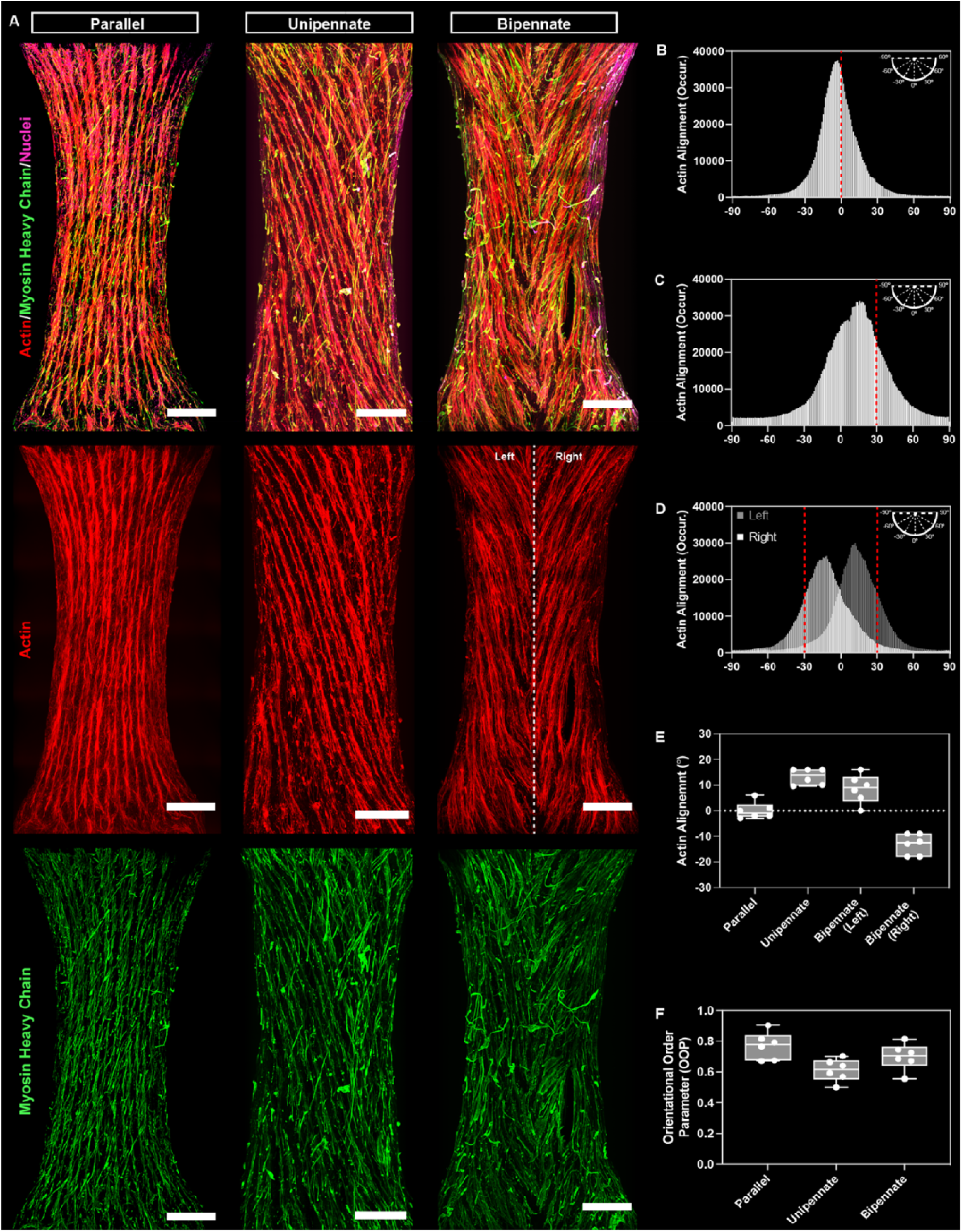
Immunofluorescent imaging the collagen scaffolds showing the ability to guide myogenesis in multiple architectures. (**A**) Immunofluorescent staining of seeded constructs after 5 days in growth media and 14 days in differentiation media, stained for actin (red), myosin heavy chain (green), and nuclei (magenta). Scale bars: 500 μm. (**B-D**) Histograms depict actin filament alignment within engineered tissues possessing (**B**) parallel, (**C**) unipennate, and (**D**) bipennate muscle architectures. An orientation angle of 0° is designated as the longitudinal axis of the scaffold. Red dashed lines indicate the intended orientation angle. (**E**) Comparison of the angle of maximum actin alignment for each construct type (n = 6 per architecture). (**F**) Comparison of the actin orientational order parameter for each construct type (n = 6 per architecture).

Quantitative analysis of the immunofluorescent images demonstrated that the vast majority of myotubes were aligned to the scaffold architectures as designed. Myotube orientation (n=6 per architecture) was consistent with the intended geometries as shown in representative histograms of actin alignment for tissues with parallel **(Fig. 3B)**, unipennate **(Fig. 3C)**, and bipennate **(Fig. 3D)** architectures. However, distribution of myotube alignment was wider than that of printed collagen filaments (Fig. 2B-D), similar to the difference between pattern and cell alignment on 2D scaffolds.^12,31^ There were also some shifts in the distribution of actin alignment (especially in the unipennate and bipennate) from the intended design towards the longitudinal axis of the engineered tissue (**Fig. 3E**). Parallel tissues were aligned to 0°, as expected, because the scaffold architecture was designed parallel to the longitudinal axis of the tissue. Unipennate and bipennate tissues, however, ranged from (+/-) 20° instead of (+/-) 30°, as designed. Cell-mediated compaction against the PDMS posts most likely caused the orientation offset towards 0°, as previous studies have shown that standard two-post designs guide myoblast alignment in the direction of tension.^30^ The orientational order parameter (OOP), where 0 is isotropic (i.e., complete lack of alignment) and 1 is complete anisotropy (i.e., perfect parallel alignment),^32^ confirmed the ability of each scaffold design to guide myotube orientation **(Fig. 3F**). Parallel exhibited the greatest anisotropy, followed by bipennate and unipennate. Mean OOP values were in the range of 0.6 to 0.8, which together with the histogram distributions (Fig. 3B to 3D) show clear alignment of the majority of the myotubes to the printed collagen scaffold architectures.

Focusing on the interaction between the collagen scaffolds and the myotubes, a tiled composite image of a representative engineered tissue with a unipennate architecture provides further insight **(Fig. 4)**. A feature that is immediately apparent (but especially at higher magnification) is the microporosity present in the printed collagen filaments (Fig. 4, inset). This results from the FRESH 3D bioprinting process, where gelatin microparticles that make up the support bath become embedded into each filament as it is extruded.^20^ Melting of the gelatin support bath and subsequent washes remove the microparticles, leaving pores behind. These pores likely facilitate a greater diffusion depth of oxygen and nutrients into the scaffold than would otherwise occur with solid filaments. Importantly, the microporosity does not prevent the scaffold’s ability to guide alignment. In fact, the cell-laden hydrogel infiltrated between the printed filaments and conformally compacted around them to yield alternating layers of collagen and fused, multinucleated myotubes.

**Fig. 4.**
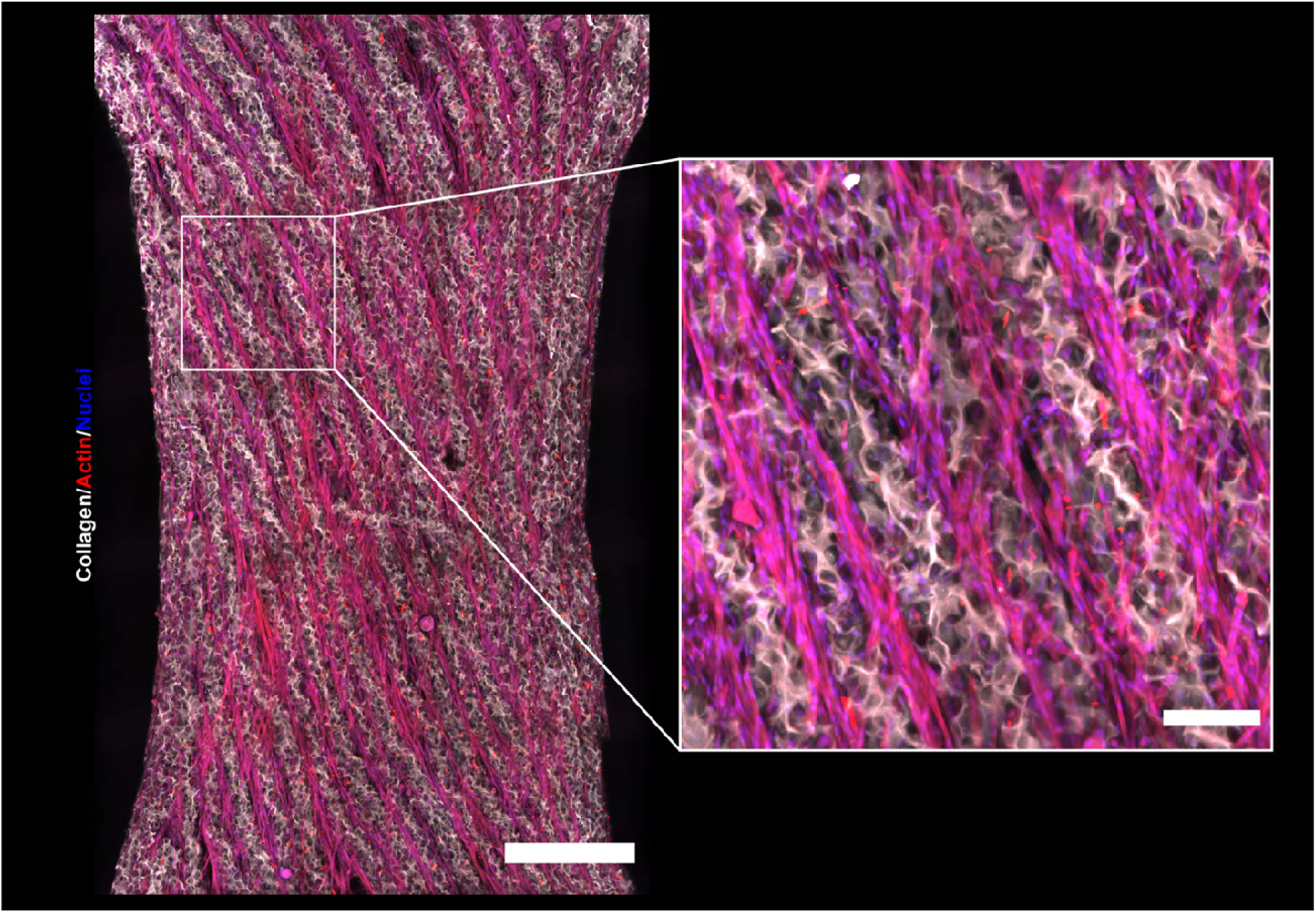
Immunofluorescent imaging of a unipennate architecture showing cellular infiltration between and conformal compaction around printed collagen filaments. The unipennate construct after 5 days in growth media followed by 14 days in differentiation media shows cellular infiltration and organization in the construct, stained for actin (red) and nuclei (magenta) with collagen reflectance (white). Scale bar: 500 μm. (**Inset**) Higher magnification image of a region of the engineered tissue revealing the spacing of collagen filaments and cell domains. Scale bar: 250 μm.

### 2.4. Calcium Handling and Force Generation in FRESH Printed Muscle Tissues

We next sought to assess the functional characteristics of the engineered muscle tissues, as the goal is to ultimately recapitulate the contractility of native tissue. Under field stimulation myotubes showed calcium transients consistent with the printed collagen scaffold architecture **(Supplementary Fig. 1A-C)**. Intracellular calcium transients for tissues with parallel, unipennate, and bipennate muscle architectures field stimulated at 1, 5, 10, and 20 Hz revealed that all tissues responded to pacing **(Supplementary Fig. 1D-F; Supplementary Videos 1-3)**. Closer examination of the calcium transient waveforms and fluorescent videos reveals that many myotubes are contracting spontaneously and independently of one another, and only synchronize when field stimulated. This suggests that most myotubes are not electrically coupled to one another, and this may be a situation where the printed collagen filaments serve as a barrier in the current scaffold design.

Contractile behavior was further investigated and quantified in terms of twitch force, summation, tetanus, and force-frequency relationship for the parallel, unipennate and bipennate architectures **(Fig. 5A-C)**. Engineered tissues (n=6 per architecture) were stimulated across a range of frequencies (1-20 Hz), and representative force traces demonstrate that twitch, summation, and tetanus occurred across all three tissue types **(Fig. 5D-F)**. These data reveal two important observations in terms of the contractile behavior. First, there was a clear trend for the twitch force to be greater in both pennate architectures as compared to the parallel tissue. This is consistent with the greater myofiber cross-sectional area in pennate muscle,^1–3^ but did not reach the level of statistical significance with our sample size. Second, contractile force as a function of stimulation frequency revealed a positive relationship for each condition **(Fig. 5G-I)**, which mimics native muscle contractile physiology.^33^ To more accurately compare the contractile properties of the engineered tissues created here to those reported in the literature, specific force (defined as twitch force divided by myotube cross-sectional area) was calculated.^34^ The specific force for parallel muscle tissues (n = 6) stimulated at 1 Hz was approximately 1 kN/m^2^. This is consistent with engineered muscle tissues created using C2C12 myoblasts, reported to have specific force values of 0.06-1.77 kN/m^2^.^24–27,34^ These calculations confirm that the myotube contractile properties are on par or greater than those reported in the literature. Twitch dynamics were also assessed, and the average twitch force traces revealed similar curves that primarily differed only in the peak force values (**Fig. 5J**). As expected from these curves, time to peak force and the time to 50% relaxation were consistent across tissues with no statistically significant differences **(Fig. 5J-L)**. This suggests that the different engineered architectures impact the twitch force of the overall construct, but not the contraction or relaxation speed of the myotubes.

**Fig. 5.**
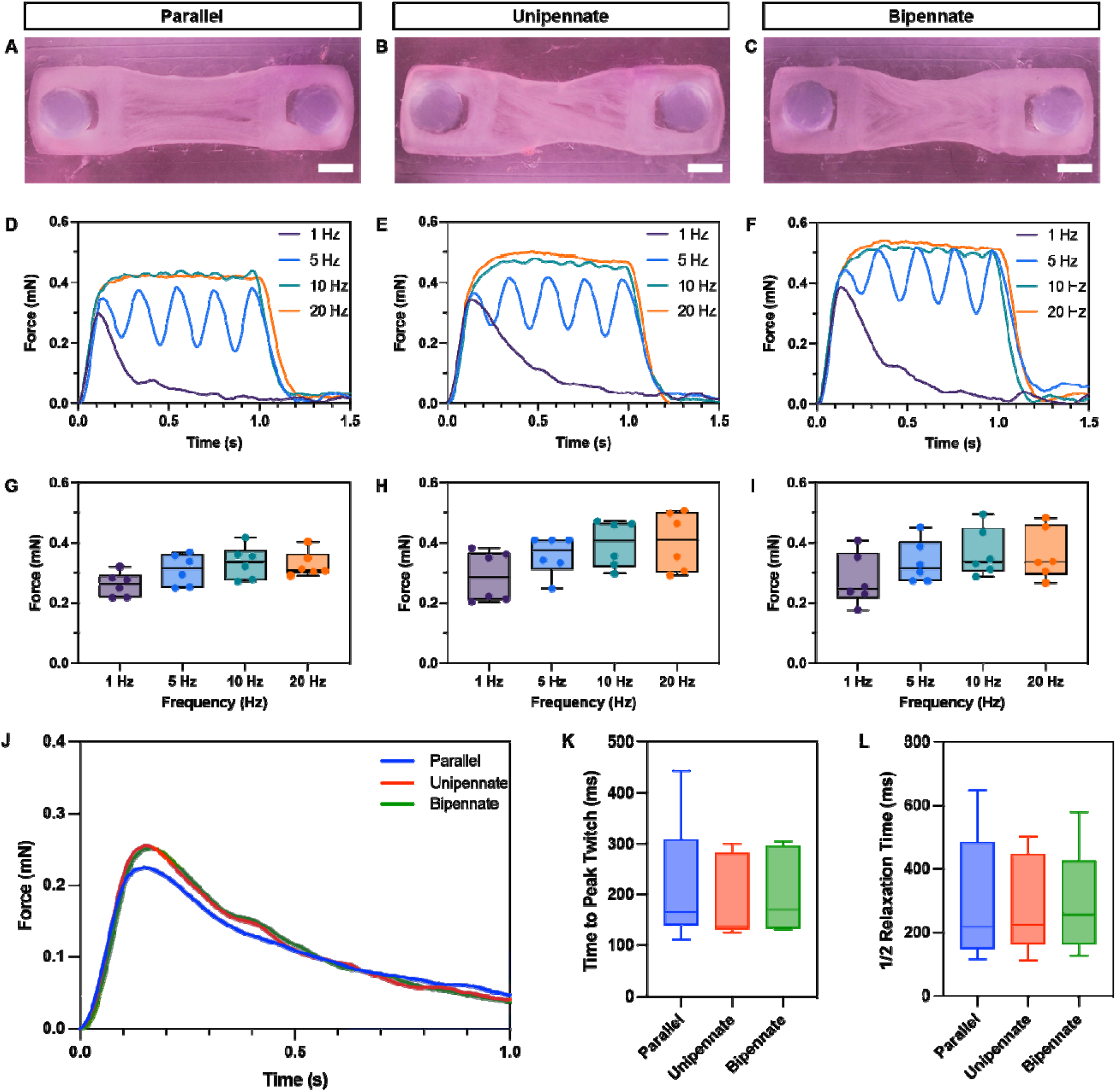
Contractile force generation of engineered muscle tissues. (**A-C**) Top-down images of engineered tissues with (**A**) parallel, (**B**) unipennate, and (**C**) bipennate architectures. (**D-F**) Tissues were subjected to field stimulation at 1, 5, 10 and 20 Hz. Representative force traces of tissues with (**D**) parallel, (**E**) unipennate, and (**F**) bipennate architectures. (**G-I**) Force-frequency plots showing contractile force as a function of stimulation frequency for (**G**) parallel, (**H**) unipennate, and (**I**) bipennate architectures. (**J**) Averaged curves for twitch force at 1 Hz showing the similar waveform morphology across tissue types. (**K**) The time to peak force generation across tissue types. (**L**) The time required for the force to relax back to ½ the peak value. All data represents n = 6 per architecture.

### 2.5. *In Vivo* Response to Engineered Scaffold Architecture

Next, we asked if the scaffold architecture, when implanted *in vivo*, could direct vascularization within the construct via collagen filament contact guidance. This is desirable because capillaries run parallel to muscle fibers *in vivo*,^35^ and we are unaware of this being demonstrated previously in the literature for engineered skeletal muscle. First, we assessed the ability of acellular collagen scaffolds to guide vascularization *in vivo* by implanting each architecture (n=4) subcutaneously in mice for 10 days **(Supplementary Fig. 2)**. Histology indicated that host cells readily invaded constructs within 10 days of implantation, which is evident by the high cell density throughout each construct. Additionally, there were some instances of vascularization, as demonstrated by the presence of red blood cells (indicated by the blue arrows). Still, there was no evidence that scaffold architecture guided the vascular ingrowth. Next, cellularized scaffolds for each architecture (n=4) were implanted into the same model to determine how pre-existing myotubes would influence the vascularization process **(Fig. 6)**. Histology indicated vascularization at the capillary scale followed the direction of scaffold architecture. Specifically, red blood cells (indicated by the blue arrows) were aligned in the direction of the printed collagen filaments and myotubes. Both parallel and unipennate architecture showed capillaries interspersed between the myotubes, all in a uniaxial direction. In contrast, the bipennate architecture had two directions of alignment, which capillaries could follow. These results are promising, suggesting that the tissues can integrate with host vasculature. Eventually, these engineered skeletal muscle tissues could potentially be a viable therapy for replacing small-scale muscle tissue defects.

**Fig. 6.**
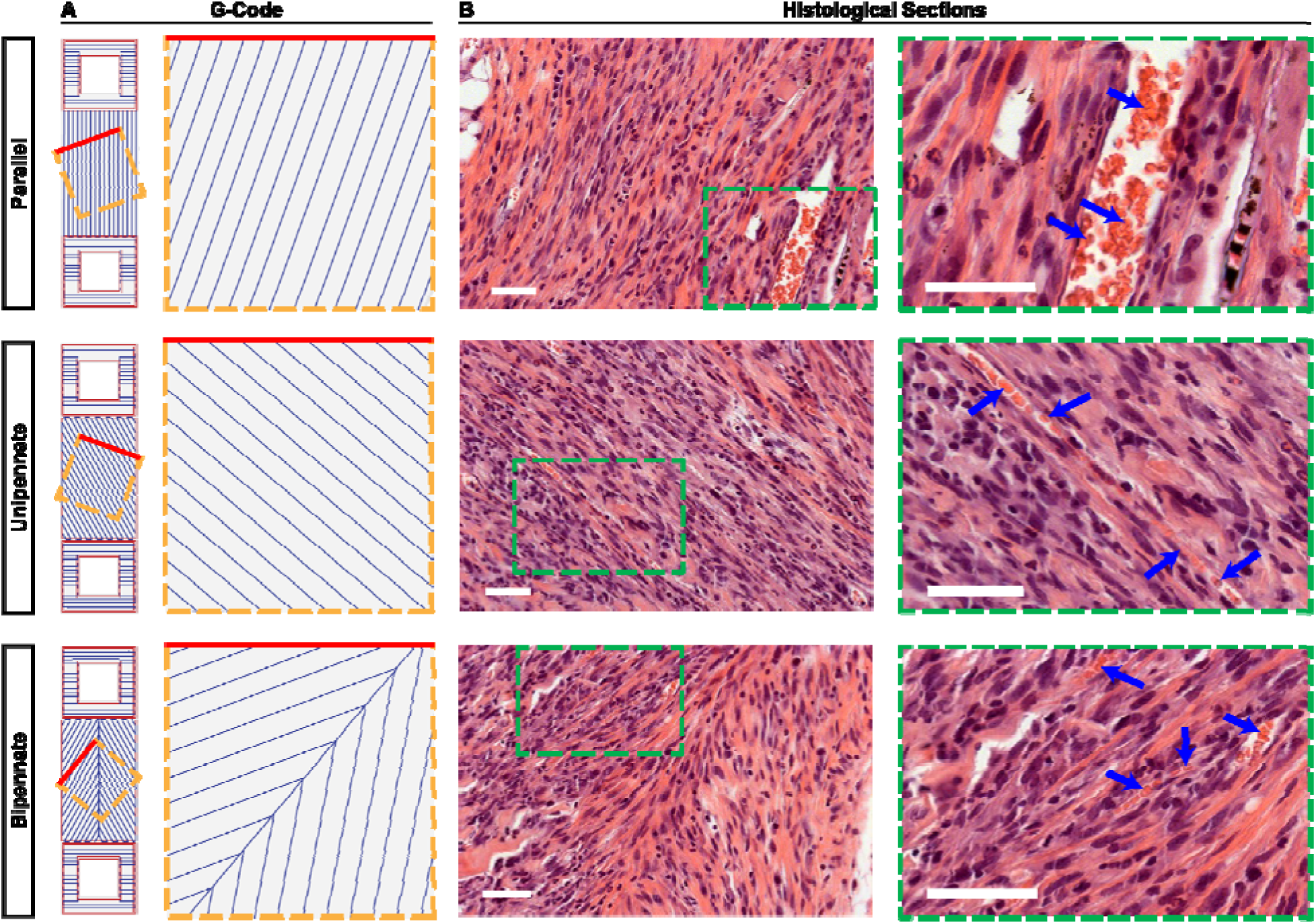
Comparison of the designed architecture and resulting cellular ingrowth and vascularization of parallel, unipennate, and bipennate tissues implanted subcutaneously in mice for 10 days. (**A**) G-code for each scaffold architecture and a zoomed in region oriented to match the scaffold architecture in the histological sections. (**B**) Histological sections stained for hematoxylin and eosin (H&E) with nuclei (purple) and cell cytoplasm/ECM (pink). The regions highlighted by the green dashed boxes indicate the areas shown at higher magnification. Blue arrows point to red blood cells, indicating host cellular infiltration and capillary formation within the muscle constructs. Scale bars: 50 μm.

### 2.6. Application to More Complex Skeletal Muscle Architectures

Finally, we extended the principles established in the initial studies to the design and engineering of more complex multipennate and convergent muscle architectures. The difference between these muscle types is the way that the fascicles, which are bundles of myofibers, are arranged within the tissue. Multipennate muscles, such as the deltoid muscle of the shoulder, consist of multiple groupings of fascicles that each insert into a single tendon, and each of these tendons taper into a common tendon. Convergent muscles, like the pectoralis major, have a broad origin where fascicles converge to a single attachment point. Here, we have designed simplified 3D models of both architectures and successfully FRESH 3D bioprinted them from collagen **(Fig. 7)**. The multipennate model, which contains multiple bipennate regions, was designed with collagen filaments aligned (+/-) 30° from the central “tendon-like” collagen section in each grouping. The OCT imaging confirmed high fidelity printing of the CAD model **(Fig. 7A)**. Engineered tissues seeded with C2C12s demonstrated alignment along the scaffold architecture, confirming that our approach can be adapted to larger, more complex muscle types **(Fig. 7B-D)**. Calcium imaging was consistent with structural imaging, showing the myotubes maintained the engineered alignment and were contractile across a range of stimulation frequencies **(Supplementary Video 4)**. The convergent model was a slightly different design challenge, because unlike all previous scaffolds where spacing between collagen filaments was constant, this varied in the convergent model. The guidance cues are wider and more spread out at one end and narrower and more tightly packed at the other end, where they converge. This required printing regions wider than a single collagen filament, but this did not impact print fidelity, which still closely matched the CAD model **(Fig. 7E)**. Engineered tissues indicated that the convergent scaffolds seeded with C2C12s were also able to control myotube alignment **(Fig. 7F-H)**. It should be noted that myotubes not contacting the collagen scaffold grew in a more random pattern, notable in the myosin signal of the convergent muscle (Fig. 7H). Calcium imaging of the convergent tissues confirmed responsiveness to electrical stimulation across a range of frequencies **(Supplementary Video 5)**. These results confirm that the methodology developed to engineer the bipennate muscle construct can be extended to engineer larger and more complex 3D muscle constructs, spanning the widest range of myofiber architectures achieved to date.

**Fig. 7.**
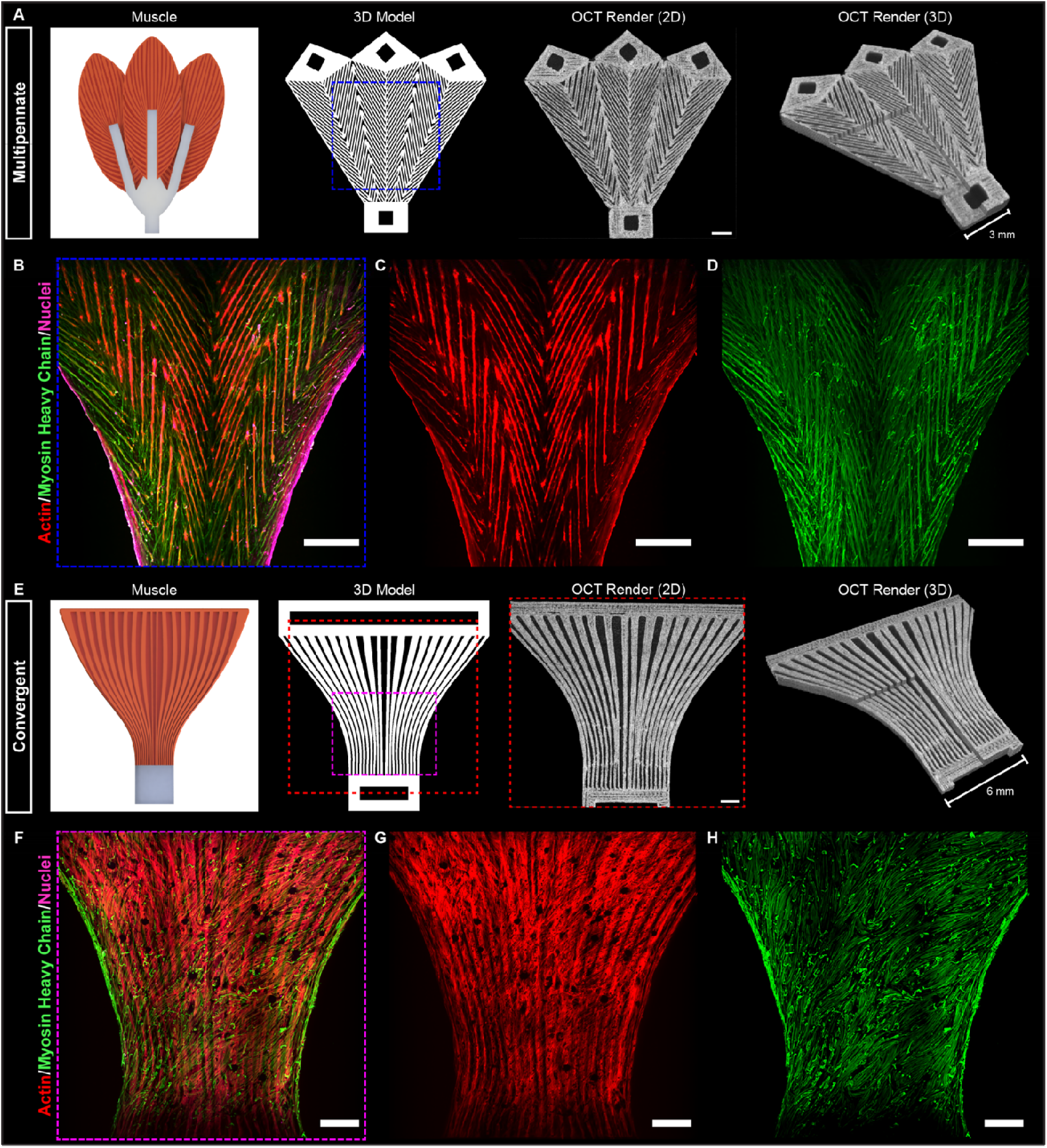
Examples of engineering larger and more complex multipennate and convergent muscle architectures. (**A**) The multipennate muscle consists of three bipennate subunits, which was converted into G-code, FRESH printed out of collagen, and structurally validated using OCT. (**B-D**) After cell seeding and culture, immunofluorescent staining of engineered tissue labeled for nuclei (magenta), actin (red), and myosin heavy chain (green). (**E**) The convergent muscle consists of variable filament spacing, which was converted into G-code, FRESH printed out of collagen, and structurally validated using OCT. (**F-H**) After cell seeding and culture, immunofluorescent staining of engineered tissues labeled for nuclei (magenta), actin (red), and myosin heavy chain (green). Scale bars: 1 mm.

## 3. Conclusions

Here we demonstrated the engineering of 3D skeletal muscle tissues with a range of complex architectures that mimic native muscle tissue. Controlling myotube alignment is a fundamental prerequisite to any tissue engineering strategy that aims to recapitulate the contractile properties of native muscle, and the FRESH 3D bioprinting platform enables this. Scaffolds mimicking a variety of muscle architectures were printed with high fidelity and proved successful in providing contact guidance to myoblasts during cell-mediated compaction, enabling differentiation into multinucleated myotubes. It is important to note that scaffold architecture alone is not sufficient to achieve muscle formation, as all scaffolds were placed on PDMS posts as anchor points to control the direction of tension during cell-mediated compaction. Further, while complex compared to other engineered muscle tissues, these constructs were still only ∼1 cm in length and ∼1 mm in thickness before compaction. These dimensions were sufficient for diffusion to maintain tissue viability, but vasculature would be required at larger scales. This is particularly a concern for therapeutic applications, where a much larger construct would be required for conditions such as VML that require multiple cubic centimeters of muscle mass. From this perspective, it is encouraging that the cellularized muscle tissues guide capillary formation *in vivo*, suggesting host-mediated vascular in-growth as a potential strategy for the vascularization of larger constructs. There, of course, are limitations, as this approach is reliant on cellular infiltration into the collagen scaffold, which depends on the motility of the seeded cells and oxygen and nutrient gradients. Although these scaffolds are relatively small, there is likely at least a small necrotic core region within each tissue due to limited oxygen and nutrient diffusion. There is also a lot of collagen present in the current scaffold designs, which clearly limits overall muscle mass in the tissue and provides a stiff element that the myotubes must contract against. This can be addressed in future work through changing the amount, design, and ECM composition of the bioink. Thus, for scaling to large tissues there is likely the need to print the muscle cells directly into the construct during the fabrication process, incorporating vascular-like channels for 3D perfusion, and tuning the bioink composition using additional ECM components, all of which have been demonstrated in FRESH printed constructs in simpler architectures and with other cell types.^20,21,36,37^ In total, these capabilities provide a clear pathway to building complex 3D muscle tissues with a range of shapes, sizes, pennation, and contractile properties for applications spanning in vitro models to *in vivo* therapies.

## 4. Methods

### Print Design of Collagen Scaffolds

Each base 3D model was created using computer-aided design (CAD) software. These models were then exported as STL files and imported into slicing software, where print parameters were selected. All collagen scaffolds were printed at 5 mm/s with 2 perimeters and a layer height of 40 µm, unless otherwise noted. Other key parameters, such as infill pattern, density, and angle were adjusted to mimic skeletal muscle architectures. For instance, the fill angle for parallel, unipennate, and bipennate skeletal muscle architectures was set to 0°, 30°, and -30°/30° from the longitudinal axis, respectively. Additionally, modifier meshes were created to apply print settings to specific regions of the model, enabling, for instance, geometric control on a layer-by-layer basis. Once the desired model was achieved, a G-code file (consisting of directions for the 3D printer) was generated.

### Bioprinter Setup

FRESH 3D bioprinting of collagen was performed on a MakerGear 3D printer modified to enable bioprinting. The thermoplastic extruder was replaced with a custom-designed syringe pump extruder, which was mounted onto the 3D printer using a custom-designed carriage. Prior to printing, collagen was prepared and loaded into a gastight Hamilton syringe (Hamilton Company). A needle (Jensen Global) with a 0.25-inch stainless steel cannula and inner diameter of 80 µm was fitted to the syringe and primed. The syringe was then mounted onto the Replistruder 4 HV.^38^

### FRESH Support Bath Generation

A gelatin microparticulate support bath (FRESH v2.0) was prepared via a complex coacervation process. Briefly, 3.0% (w/v) gelatin type B (Fisher Chemical), 0.125% (w/v) Pluronic® F-127 (Sigma-Aldrich), and 0.25% (w/v) gum Arabic (Sigma-Aldrich) were dissolved in a 1 L beaker containing a 50% (v/v) ethanol solution heated to 45°C. The solution was mixed on a stir plate until homogeneous, and pH was subsequently adjusted to 5.35-5.65 (depending on desired particle size) through the dropwise addition of 2 N hydrochloric acid (HCl). The beaker was then transferred to a temperature-controlled room, where it was placed under an overhead stirrer and sealed with Parafilm to minimize evaporation. The solution was allowed to cool to room temperature overnight as it was stirred at approximately 500 RPM to form the gelatin microparticles. The following day, the microparticles were allowed to settle, and the supernatant was poured off. The resulting slurry was transferred to 250-mL Nalgene centrifuge bottles, where it was compacted via centrifugation at 500 G for 3 minutes. The supernatant was poured off, and the gelatin was resuspended in deionized water and centrifuged at 750 G for 3 minutes to wash away remnants of ethanol and Pluronic® F-127. Washes in deionized water were repeated an additional two more times (at 750 G and 1000 G for five minutes each) for three total washes. Slurry was then washed three times in 50mM HEPES, pH 7.4 buffer solution. Uncompacted gelatin slurry in the buffer solution was stored at 4°C for up to one month. Prior to printing, uncompacted slurry was placed in a vacuum chamber for 15-30 minutes and then centrifuged at 1000 G for 5 minutes. Supernatant was poured off, and the compacted slurry was then transferred to a small Petri dish for printing.

### FRESH 3D Bioprinting of Type I Collagen

3D bioprinting of type I collagen was conducted using a modified open-source 3D bioprinter. Acidified collagen at approximately 24 mg/mL (LifeInk^®^ 200, Advanced BioMatrix, diluted with 0.24 M acetic) was loaded into a 2.5-mL gastight Hamilton syringe (Hamilton Company) and fitted with a 34-gauge needle (Jensen Global) with a 0.25-inch stainless steel cannula and inner diameter of 80 µm, unless noted otherwise. The syringe was then loaded into the Replistruder 4 and primed. Compacted FRESH support bath was transferred to an adequately sized print container and secured to the print bed with double-sided tape. The needle was manually positioned above the center of the print container and lowered into the slurry, leaving approximately 1 mm from the bottom of the container. The print was started using a Duet web control interface. Upon print completion, the print container was transferred to a 37°C oven, where the gelatin slowly melted, releasing the printed collagen construct. Gelatin was rinsed away with 3 or more successive washes with warm 50 mM HEPES, pH 7.4 buffer solution supplemented with 1% (v/v) Penicillin-Streptomycin (10,000 units/mL penicillin, 10,000 µg/mL streptomycin) (15140-122, Life Technologies). Final prints were sterilized under UV Ozone for 15 minutes. Print containers were wrapped in Parafilm and stored at 4°C.

### Optical Coherence Tomography of FRESH 3D Bioprinted Collagen Constructs

Large volume imaging of collagen constructs was conducted using a 1300 nm optical coherence tomography (OCT) system (VEG210C1, ThorLabs). This system is equipped with a lens kit with 13 μm lateral resolution and has an imaging depth of 11 mm. Image processing was performed using ImageJ (https://imagej.net/Welcome).

### Cell Culture

C2C12 myoblasts (CRL-1772, ATCC) were cultured at 37°C under 10% CO_2_ in growth media consisting of Dulbecco’s Modified Eagle Medium (DMEM) – high glucose (15-013-CM, Corning) supplemented with 1% (v/v) Penicillin-Streptomycin (10,000 units/mL penicillin, 10,000 µg/mL streptomycin) (15140-122, Life Technologies); 1% (v/v) L-Glutamine (200 mM) (25030-081, Life Technologies); and 10% (v/v) Fetal Bovine Serum (FBS) (89510-186, VWR). Media was exchanged every two days, and cells were passaged upon reaching 80% confluence. On the day 5 following cell seeding on the collagen constructs, the culture media was switched to differentiation media consisting of DMEM – high glucose (15-013-CM, Corning) supplemented with 1% (v/v) Penicillin-Streptomycin (10,000 units/mL penicillin, 10,000 µg/mL streptomycin) (15140-122, Life Technologies); 1% (v/v) L-Glutamine (200 mM) (25030-081, Life Technologies); and 2% (v/v) Horse Serum (HS) (45001-058, VWR). Tissues were maintained in differentiation media for 2 weeks with media exchanges performed every 2 days.

### PDMS Chamber Fabrication

Custom-designed polydimethylsiloxane (PDMS) chambers were created to house the collagen constructs during casting and culture. Each chamber consisted of a rectangular well with two cylindrical posts at either end that served as anchor points for the tissue. To create each mold, the negative mold was first designed in CAD software and then printed out of polylactic acid (PLA) using a Prusa 3D printer. Negative molds were then placed into large Petri dishes. Sylgard 184 PDMS prepolymer (Dow Corning) was prepared by mixing the base and curing agent in a 10:1 weight ratio in a Thinky Conditioning Mixer (Phoenix Equipment, Inc.) using a cycle of 2 minutes of mixing at 2000 G followed by 2 minutes of degassing at 2000 G. The prepolymer was cast into the molds and placed in a vacuum chamber for 60 minutes to remove any air bubbles. PDMS chambers were then cured overnight in a 65°C oven. After curing was complete, the chambers were carefully removed from the negative molds and placed in beakers to be cleaned. The cleaning and disinfecting regimen consisted of three cycles in a sonicating bath: (i) sonicate in a beaker of 1% Enzol for 30 minutes; (ii) wash with DI water and sonicate in a beaker of DI water for 30 minutes; (iii) wash with DI water and sonicate in a beaker of 70% ethanol for 60 minutes. Chambers were then dried in the biosafety cabinet prior to conformal casting of collagen constructs.

### Conformal Casting of Collagen Constructs

Sterile collagen scaffolds stored in 50 mM HEPES supplemented with 1% (v/v) Penicillin-Streptomycin (10,000 units/mL penicillin, 10,000 µg/mL streptomycin) (15140-122, Life Technologies) were transferred to the biosafety cabinet. Vacuum grease was sterilized under UV Ozone for 15 minutes. PDMS chambers were secured to the bottom of a well in a 12-well plate with vacuum grease (BD Corning). Each PDMS well was incubated with 1% (w/v) Pluronic F-127 (Sigma) for 45 minutes to prevent cell attachment to the PDMS well interior. The Pluronic F-127 solution was then aspirated, and PDMS wells were rinsed with sterile 1x PBS three times. Fresh 50 mM HEPES was added to each well to prevent collagen scaffolds from drying out during placement. Collagen scaffolds were then carefully transferred to the chambers with a sterile surgical hook, where they were anchored around the two PDMS posts using sterile forceps. Immediately prior to cell casting, 50 mM HEPES was aspirated from all PDMS wells. Cells were then cast around the collagen scaffolds inside the PDMS wells at a final concentration of 30 million cells/mL in solution of neutralized collagen (0.5 mg/mL) and 10% v/v Matrigel (Corning). To do this, the cell suspension was added to a microcentrifuge tube containing collagen (10 mg/mL), 2.3% v/v 1 N NaOH, 5% v/v 10x PBS, and 10% v/v Matrigel and quickly mixed; a 70-µL aliquot was then pipetted into each well. Care was taken to prevent the introduction of any bubbles to the cell suspension. Scaffolds were then incubated at 37°C for 60 minutes to allow collagen gel formation, after which 3 mL of warm growth media were added to each well. All casted constructs were initially cultured in growth media, with media exchanges every 2 days. On day 5 after seeding, culture media was switched to differentiation media and cultured for up to 14 days, with media changes every 2 days.

### Immunofluorescence Staining and Imaging

Skeletal muscle tissues were fixed in a solution comprised of 4% formaldehyde (15710, Electron Microscopy Sciences) in 1x DPBS. Triton X-100 (ThermoFisher Scientific) was added at 1:200 to aid in cell permeabilization. Tissues were fixed in the solution for 1 hour and subsequently rinsed with 1x PBS three times to remove any formaldehyde. Tissues were blocked with a 5% goat serum solution in 1x PBS overnight and washed with 1x PBS three times. Tissues were then stained with mouse anti-myosin heavy chain (MHC) (MA5-11748, ThermoFisher Scientific) at a dilution of 1:100 in 1x PBS through incubation with the primary antibody at 4°C for 2 days on a rotary shaker. Tissues were washed with 1x PBS three times and then stained with goat anti-mouse secondary antibody conjugated to AlexFluor488 (A28175, Life Technologies) at 1:400 dilution, TO-PRO-3 stain (T3605, ThermoFisher Scientific) at a 1:400 dilution to stain for nuclei, and phalloidin conjugated to AlexaFluor 555 (A22284, Life Technologies) at a 3:200 dilution to stain for actin. Tissues were incubated for an additional 2 days on a rotary shaker at 4°C and washed three times in PBS prior to imaging. To increase the imaging penetration depth and decrease light scattering, representative engineered tissues were cleared by successive dehydration steps in isopropanol (incubation in solutions of 25% v/v, 50% v/v, 75% v/v, and 100% v/v IPA in 1x PBS for 30 minutes each) and subsequent clearing via incubation in a solution of benzyl alcohol (Sigma-Aldrich) and benzyl benzoate (Sigma-Aldrich) at a 1:2 ratio. A Nikon A1R MP+ multiphoton confocal microscope and 16x (0.80 NA) objective were used to obtain tile scans and 3D z-stacks of all tissues.

### Cellular Alignment Analysis

Maximum intensity projections of Z-stacks were used to quantify actin alignment in engineered tissues (n = 6 per muscle architecture, about 150 μm in depth). The angular distribution of actin filaments was analyzed using a custom MATLAB code, which has previously been used in cellular alignment quantification and is based off a ridge detection algorithm used for fingerprint enhancement.^32,39^ This was executed by applying a threshold to the actin channel, which enables the creation of a binary mask of actin location that is then used to generate orientation vector maps. Histograms were created of actin orientation angles, and a 2D orientation order parameter (OOP) was calculated for each tissue, which is a metric that describes the degree of alignment, where 0 corresponds to no alignment (random orientation of actin filaments) and 1 corresponds to perfect alignment (anisotropic orientation of coaligned filaments).

### Calcium Imaging of Muscle Tissues

Electrophysiology of skeletal muscle tissues was assessed via calcium imaging during spontaneous and stimulated muscle contractions. Tissues were cultured for approximately 14 days before imaging. Each construct was first transferred to a small dish and rinsed with Tyrode’s solution to remove excess media. Tissues were then incubated in Tyrode’s solution with 5 µM calcium indicator Cal 520 AM (21130, AAT Bioquest) and 0.25% Pluronic F-127 (P2443, Sigma Aldrich) for 60 minutes at 37°C, followed by a 30-minute incubation at room temperature. Tissues were then washed with Tyrode’s solution and placed in a small petri dish filled with solution for imaging. The dish was loaded into a heated stage maintained at 37°C, and an imaging platform consisting of an epifluorescent stereomicroscope (SMZ1000, Nikon), GFP filter, X-Cite lamp (Exceltias), and Prime 95B Scientific CMOS camera (Photometrics) was used for high-speed imaging of calcium transients at a frame rate of 100 frames per second. Field stimulation was achieved by placing two parallel platinum electrodes into the dish and using a Grass Stimulator to apply a square wave pulse at a range of frequencies from 1 to 20 Hz (1, 5, 10, and 20 Hz) and 90 V for 10 ms.

### Measurement of Tissue Contractile Forces

Skeletal muscle tissues were cultured for 14 days prior to force measurements. To conduct measurements, the PDMS mold housing the tissue was first removed from its 12-well plate using sterile forceps and transferred to a 6-well plate. Using forceps, the tissue was then transferred to another Petri dish containing a mounted stainless steel pin and fresh Tyrode’s solution. This dish was then loaded into a heated stage maintained at 37°C. The stainless steel pin was adjusted to be in line with a custom-mounted optical force transducer (World Precision Instruments). At one end, the open end of the tissue was carefully placed on the stainless steel pin in the Petri dish. The other end was placed around another stainless steel pin attached to the transducer. The transducer was situated on linear actuators, which were used to position the transducer at a distance from the opposing stainless steel pin equal to the resting length of the tissue (L_0_). The heated stage was then placed under a stereomicroscope. A DSLR camera mounted on the stereomicroscope was used to record tissue elongation during force measurements, and a custom LabVIEW program was used to record tissue forces. A Grass Stimulator was used to stimulate the tissue and induce contractions by delivering a 90 V square wave pulse at 1 Hz. The tissue was then stretched using the linear actuators in ∼2% increments and stimulated at each. At the final length (corresponding to ∼20% elongation), the tissue was stimulated at 1, 5, 10, and 20 Hz. Force traces were recorded at each. Force as a function of time for each frequency was plotted to demonstrate force-frequency relationships. Contractile forces were determined by subtracting baseline force from peak force amplitudes.

### Animal Studies

Animal care and experimental procedures were carried out at Carnegie Mellon University in accordance with the National Institutes of Health Guide for the Care and Use of Laboratory Animals under an approved Institutional Animal Care and Use Committee (IACUC) protocol. C57BL/6 male mice (6 to 8 weeks old; 22-26 grams) were utilized in this study. Scaffolds were implanted subcutaneously on right side of the mice dorsum under general anesthesia (2% isoflurane). Constructs were harvested 10 days post-implantation to evaluate vascular ingrowth. At harvest, scaffolds were cleaned of connective tissues, adipose tissue and immediately fixed in 10% formalin overnight, and then embedded in paraffin. Embedded tissues were then sectioned and stained with hematoxylin and eosin (H&E) to visualize cellular ingrowth.

## Acknowledgements

This research was supported by the Additional Ventures Foundation Cures Collaborative, the National Science Foundation Graduate Research Fellowship Program under Grant No. DGE 1745016, and the Army Research Office under Cooperative Agreement Number W911NF-23-2-0138. Any opinions, findings, and conclusions or recommendations expressed in this material are those of the authors and do not necessarily reflect the views of the National Science Foundation. The views and conclusions contained in this document are those of the authors and should not be interpreted as representing the official policies, either expressed or implied, of the Army Research Office or the U.S. Government. The U.S. Government is authorized to reproduce and distribute reprints for Government purposes notwithstanding any copyright notation herein.

