## Supplementary Materials for "Engineering 3D Skeletal Muscle Tissue with Complex Multipennate Myofiber Architectures"

### SUPPLEMENTARY VIDEOS

**Supplementary Video 1. Field stimulation of an engineered skeletal muscle tissue with a parallel architecture.** A representative tissue was stained with a calcium-sensitive dye and subjected to field stimulation at 1, 5, 10, and 20 Hz. The tissue responds to pacing and appears to approach tetanus at higher frequencies. Scale bar: 1 mm.

**Supplementary Video 2. Field stimulation of an engineered skeletal muscle tissue with a unipennate architecture.** A representative tissue was stained with a calcium-sensitive dye and subjected to field stimulation at 1, 5, 10, and 20 Hz. The tissue responds to pacing and appears to approach tetanus at higher frequencies. Scale bar: 1 mm.

**Supplementary Video 3. Field stimulation of an engineered skeletal muscle tissue with a bipennate architecture.** A representative tissue was stained with a calcium-sensitive dye and subjected to field stimulation at 1, 5, 10, and 20 Hz. The tissue responds to pacing and appears to approach tetanus at higher frequencies. Scale bar: 1 mm.

**Supplementary Video 4. Field stimulation of an engineered skeletal muscle tissue with a multipennate architecture.** A representative tissue was stained with a calcium-sensitive dye and subjected to field stimulation at 1, 5, 10, and 20 Hz. The tissue responds to pacing and appears to approach tetanus at higher frequencies. Scale bar: 1 mm.

**Supplementary Video 5. Field stimulation of an engineered skeletal muscle tissue with a convergent architecture.** A representative tissue was stained with a calcium-sensitive dye and subjected to field stimulation at 1, 5, 10, and 20 Hz. The tissue responds to pacing and appears to approach tetanus at higher frequencies. Scale bar: 1 mm.

### SUPPLEMENTARY FIGURES

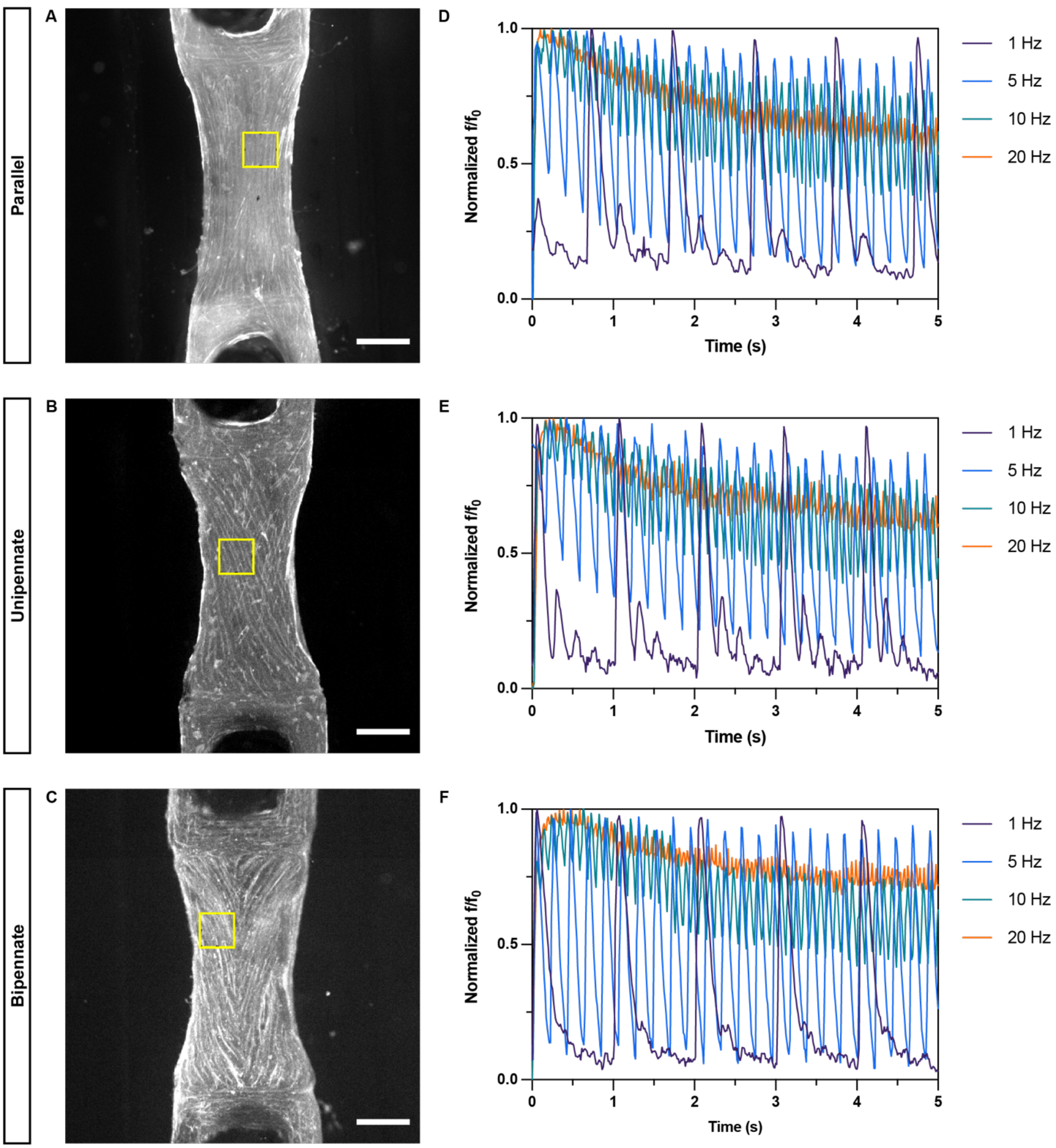


**Fig. S1. Calcium imaging demonstrates contractile myotubes throughout engineered muscle tissues.** Top view of representative engineered muscle tissues stained with a calcium-sensitive dye, revealing the underlying (**A**) parallel, (**B**) unipennate, and (**C**) bipennate muscle architectures. Scale bars: 1 mm. Tissues were field stimulated over a range of frequencies (1-20 Hz). Traces of intracellular calcium signal during field stimulation were optically recorded for tissues possessing (**D**) parallel, (**E**) unipennate, and (**F**) bipennate muscle architectures. Calcium fluorescence intensity for each sample was normalized over a range from 0 to 1, where 0 indicates the baseline relaxed state and 1 indicates the peak measured calcium level.

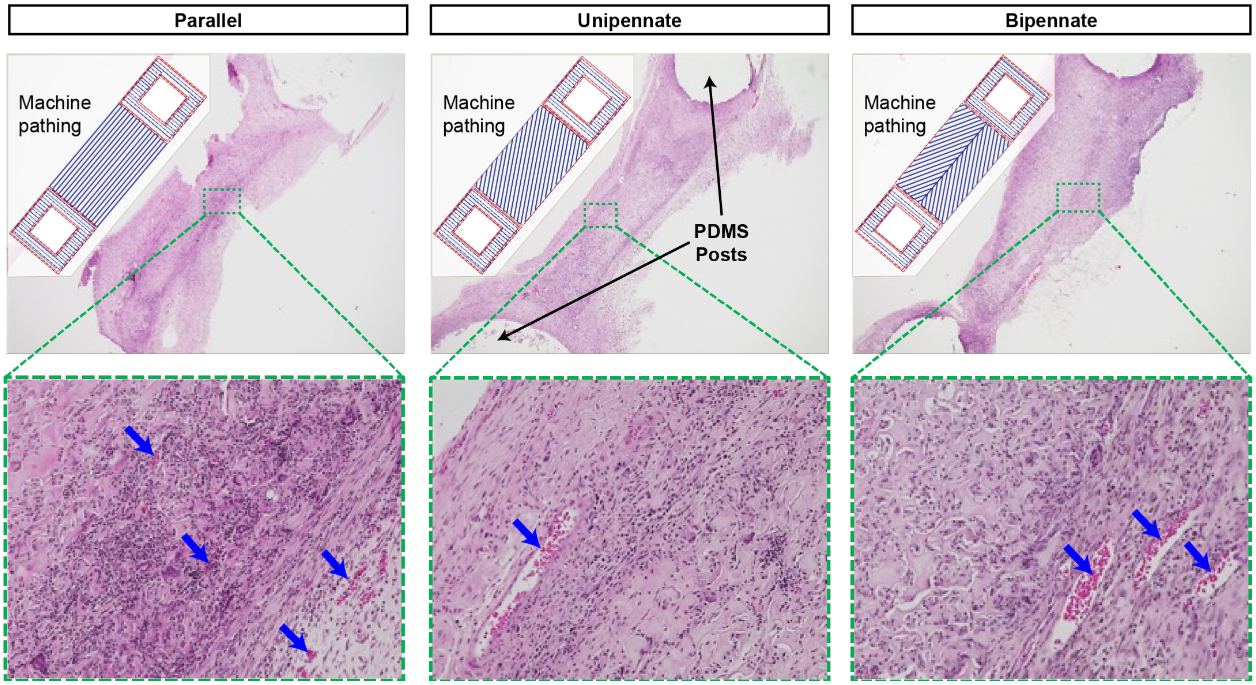


**Fig. S2. Histological sections of acellular collagen scaffolds with parallel, unipennate, and bipennate architectures implanted subcutaneously in mice for 10 days.** Hematoxylin and eosin (H&E) staining for nuclei (dark purple) and cell cytoplasm/ECM (pink). Blue arrows point to red blood cells, indicating host cellular infiltration.
